# The longevity of cut flowers of Iranian narcissus is affected by genotype, physiochemical characteristics and stem structural factors

**DOI:** 10.1101/2024.08.16.608278

**Authors:** Azra Haghshenas, Abolfazl Jowkar

## Abstract

Flower longevity is one of the critical factors in determining the quality of cut flowers. narcissus, a popular cut flower in Iran, is cultivated outdoors in the fall and winter. Despite numerous studies on extending the longevity of narcissus flowers, our understanding of internal changes in floral tissues during the postharvest period is limited. This study aimed to investigate the morphophysiological and biochemical changes in floral tissues during postharvest conditions in Iranian narcissus populations. Ten narcissus populations were collected from various regions of Iran and cultivated in a greenhouse. The harvested narcissus flowers were kept in vases containing distilled water. Flower longevity; physicochemical characteristics, such as relative solution uptake, electrical conductivity, relative water content, total protein, antioxidant enzyme activity, carotenoids, H_2_O_2_ and malondialdehyde; stem mucilage sugars; structural features of the stem and vascular bundles; and the number of bacteria in the vase solution were evaluated on the first and third days after harvest. The experiment was conducted in a completely randomized design with nine replications, and mean comparisons were performed via SPSS software. The longevity of flowers is influenced by genetic characteristics and floral tissues. The populations of Shahla from the Khusf region and Porpar from the Khafr region presented the greatest longevity, whereas white narcissus presented the shortest longevity. Compared with petal tissue, corona tissue has greater longevity because of its thicker structure, higher water retention capacity, lower production of free radicals, and stronger antioxidant defense system. The number of vascular bundles was significantly positively correlated with relative solution uptake. No vascular blockage was observed during the postharvest period, but stem cells collapsed on the third day of vase life, which seemed to affect water uptake. The reason for the lack of vascular blockage in the narcissus stem requires further research.

## 1 Introduction

One of the critical attributes determining the quality of cut flowers is their longevity and postharvest life (Naing et al., 2022), which depend on genetic factors, preharvest conditions, and both harvesting and postharvest conditions (Verdonk et al., 2023; Rabiza-Świder et al., 2020b; Vijayakumar et al., 2021). Extending the longevity of flowers has always been a challenge in the floriculture industry, both in local and global markets (Naing et al., 2022). More durable flowers not only have greater market appeal (Naing et al., 2022; Cavallaro et al., 2023) but also help reduce resource waste and increase producer profits. Among cut flowers, narcissus is highly popular worldwide, especially in Iran, owing to its unique appearance, color, and pleasant fragrance. It is classified as a specialty cut flower (SCF), which is typically produced outdoors during specific seasons (Darras, 2021). In Iran, on the basis of local observations, narcissus blooms in autumn and winter, growing both in natural habitats and in cultivated fields and gardens under fruit trees, and is sold as a cut flower. Despite numerous postharvest studies on narcissus, our understanding of the physicochemical and structural factors affecting the aging process and longevity of cut narcissus flowers remains insufficient.

Aging is a complex process that leads to the death of an organ or the entire organism. At the cellular level, it involves a series of events, such as changes in membrane permeability, degradation of proteins and nucleic acids, alterations in sugar status, and the activity of antioxidant enzymes such as peroxidases, superoxide dismutase, and catalase. The transfer of nutrients from aging organs to other parts of the plant results in programmed cell death (PCD) and eventually leads to the death of the organ or the entire organism (Gul et al., 2020). Kondo et al. (2020) reported that aging occurs in both cut carnations and those still attached to the mother plant; however, in cut flowers, the rapid decline in sugar levels accelerates ethylene production, leading to faster aging. While the color, fragrance, and shape of flowers increase their aesthetic appeal and market value, they also play crucial roles in attracting pollinators and completing reproductive processes, which contributes to plant survival. Once pollination occurs, plants no longer need to invest energy in maintaining attractive flower parts such as petals, leading to their eventual senescence (Sun et al., 2021; Gul et al., 2020). The limited longevity of flowers is due primarily to the role these organs play in the plant lifecycle; however, under postharvest conditions, longevity, in addition to genetic traits, is influenced by factors such as water stress, environmental humidity, temperature, ethylene levels, nutrient deficiencies for respiration, vascular occlusion, microbial activity, and many other factors (Gul et al., 2020).

Postharvest changes at the end of a cut stem can affect water uptake, thereby influencing a plant’s physiochemical processes and longevity. One significant change is vascular blockage at the cut end of the stem, which can be caused by physiological factors such as tylosis formation and programmed cell death, microbial factors such as bacteria, physical factors such as air bubble formation in xylem vessels, and the deposition of gum and mucilage in vessels (Da Silva, 2015). The cut flowers continuously lose water through evaporation and transpiration, and if they cannot compensate for this loss due to vascular blockage, they soon experience water stress (Manzoor et al., 2024; Da Silva, 2015), leading to premature aging and reduced longevity of the cut flower (Da Silva, 2015). According to the current literature, no studies have specifically investigated changes in stem tissue and vascular bundles at the cut end of narcissus, particularly the Iranian varieties.

Numerous studies have investigated treatments to extend the vase life of narcissus flowers. However, our understanding of the morphological changes in stem tissues, the internal physiological and biochemical factors in the corona and petal tissues affecting vase life, and how these factors contribute to a better comprehension of the senescence process in narcissus is still limited. This study examines the vase life of ten Iranian narcissus populations with the aim of investigating the impact of physiological and biochemical changes in corona and petal tissues, as well as alterations in stem tissue during the postharvest period. Understanding the trends in morphological, physiological, and biochemical changes within the inflorescence will not only facilitate the selection of treatments that increase vase life but also contribute to the breeding of flowers with improved durability, thereby reducing waste and increasing producer revenue.

## 2 Materials and methods

## 2-1 Plant material and growth conditions

Bulbs from ten populations of Iranian narcissus, including *Narcissus tazetta* and *Narcissus papyraceus* (commonly known as white narcissus), were collected from natural habitats, old farms and gardens across various geographical regions of Iran (Haghshenas et al., 2024). The collection included one white population and one Shahla population from Shiraz; one Shahla population from Kazerun; one Porpar population and one Shahla population from Khafr; one Shahla population from Abdanan; one Sadpar population and one Shahla population from Behbahan; one Shahla population from Juybar; and one Shahla population from Khusf. The bulbs were planted in early autumn in the greenhouse of the Horticultural Sciences Department at Shiraz University (35°52′N and 29°43′E) in soil with a 4:1 ratio of silty-clay soil and leaf mold. They were planted in rows spaced 20 cm apart, with bulbs 18 cm apart within rows. Irrigation was carried out when the soil dried, the greenhouse temperature during the growing season was maintained at 23±5°C, and natural light was used.

## 2-2 Harvesting and Sample Preparation

The flowers were harvested when the first flower on the inflorescence just stated opening and were immediately transferred to the laboratory. The stems were trimmed to a length of 10 cm under water with a sharp blade and placed in 300 ml plastic cups filled with 250 ml of distilled water, with the cups covered with plastic wrap. To assess flower longevity, the flower cups were kept in a growth chamber (Noor Sanat Company, Iran) at 23°C with 60% relative humidity, a light intensity of 10 μmol/m²s, and a light/dark cycle of 12 hours. The same plastic cups used as flower vases were also used as controls; they were filled with 250 millilitres of distilled water, covered with plastic wrap with a hole in the center, and placed in the same growth chamber alongside the flowers.

## 2-3 Vase Life

The visual characteristics of the flowers, such as the number of opened flowers, color changes, and wilting of the petals and corona, were monitored and recorded daily. The day of harvest was considered day 0, and the duration until 50% of the flowers in the inflorescence wilted and the inflorescence lost its freshness and beauty was considered the vase life. To assess vase life, nine stems of narcissus flowers were used. Additionally, the lifespans of petals and coronas were considered separately from the lifespan of the inflorescence (Rabiza-Świder et al., 2020b). When the petal edges turned brown, it was recorded as the end of the petal life, and the day the corona wilted and lost its turgidity was recorded as the end of the corona life.

## 2-4 Measurement of Morphophysiological Characteristics

At the beginning of the experiment, the number of buds on each inflorescence was counted, and the cut flowers were weighed. To calculate water uptake, relative water uptake, and evaporation and transpiration rates, the weights of the vase containing water and flowers, as well as the weights of the water and control vases, were measured daily via a digital scale with a precision of two decimal places. Additionally, the dry weight, moisture content, and dry matter or biomass of the corona and petal tissues were separately calculated. The stem diameter was measured with a digital caliper.

## 2-4-1 Water Uptake (WU) and Relative Water Uptake (RWU)

The amount of evapotranspiration was calculated on the basis of the difference in the weights of the vases at the start and end of the experiment. Relative water uptake (RWU) was determined via a slight modification of the method of Spricigo et al. (2021) by dividing the amount of solution absorbed by the initial weight of the flowers (Spricigo et al., 2021). The amount of water uptake (WU) was calculated by the difference in the weight of the vase with flowers on the initial day and the final day minus the evaporation rate. Evaporation was calculated as the difference in the weight of the control vase between the start and end of the experiment:

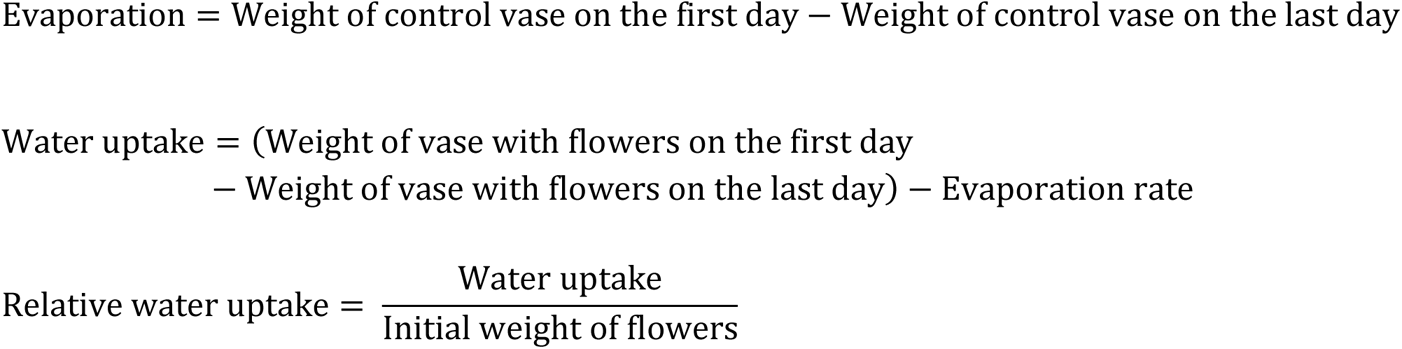

## 2-4-2 Water Content

The relative water content (RWC) of the tissues in the corona and petals was measured in nine replicates on the day of harvest and on the third day. The petals and corona were carefully and precisely separated and weighed using a digital scale (wet weight). The flower tissues were subsequently placed in Petri dishes and covered with distilled water for 24 hours. After this, the tissues were placed on paper towels for 5 minutes to remove excess water and weighed again (turgid weight). The samples were then placed in an oven at 70°C for 48 hours and weighed once more (dry weight). The RWC was calculated via the following formula (with slight modifications to the method of Rabiza-Świder et al., 2020b):

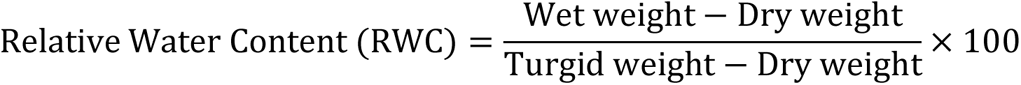

## 2-4-3 Ion Leakage and Electrical Conductivity

To measure cellular ion leakage, ten 5 mm diameter discs of the corona and petal tissues were separated on the day of harvest and on the third day. The discs were placed in 10 ml of double-distilled water and kept on a shaker at room temperature for 24 hours. Electrical conductivity was measured via a Metrohm 644 conductometer (Switzerland), and pH was measured via a Jenway pH meter (UK). Electrical conductivity was reported in μS/cm (Rabiza-Świder et al., 2020b).

## 2-4-4 Examination of Cross-Sectional Stem Characteristics via Microscopy Image Processing

To study the stem structure and its changes during the postharvest period of cut narcissus flowers, cross-sections of stems that were kept in floral preservative solution for three days were compared with cross-sections of freshly harvested stems. For each population, nine stems that were freshly harvested and nine stems that had been kept in the floral preservative for three days were used. For sample preparation, 0.5 cm segments from the ends of freshly harvested stems and stems from the third day were cut and fixed in FAA solution (formalin 5 ml, acetic acid 5 ml, and 90 ml 50% ethanol; Lear et al., 2022). The thin sections were subsequently sliced by hand via a sharp blade. The samples were immediately placed on slides and observed under a polarized microscope at magnifications of 25x (Olympus BH2, Japan) and 100x (Olympus BH41, Japan), and images were captured. Three sections per sample were examined under a microscope. Microscopy images were taken with a mobile phone camera (Sony Xperia Z5, Japan), and quantitative features of the stem and vascular bundles were calculated via ImageJ software version 15.3c (Figure 1). These features include:

- Stem perimeter
- Stem area/total cross-sectional area
- Inner stem thickness
- Hollow cavity area of the stem
- Min Feret index of the stem
- Max Feret index of the stem
- Solidity index of the stem
- Stem circularity index
- Minor indices of the stem
- Major indices of the stem
- Vascular bundle number per stem
- Vascular stem area
- Vascular bundle perimeter
- Vascular circularity index
- Min Feret index of vascular bundles
- Max Feret index of vascular bundles
- Minor indices of vascular bundles
- Major indices of vascular bundles
- Ratio of the vascular tissue area to the stem area

**Fig. 1.**
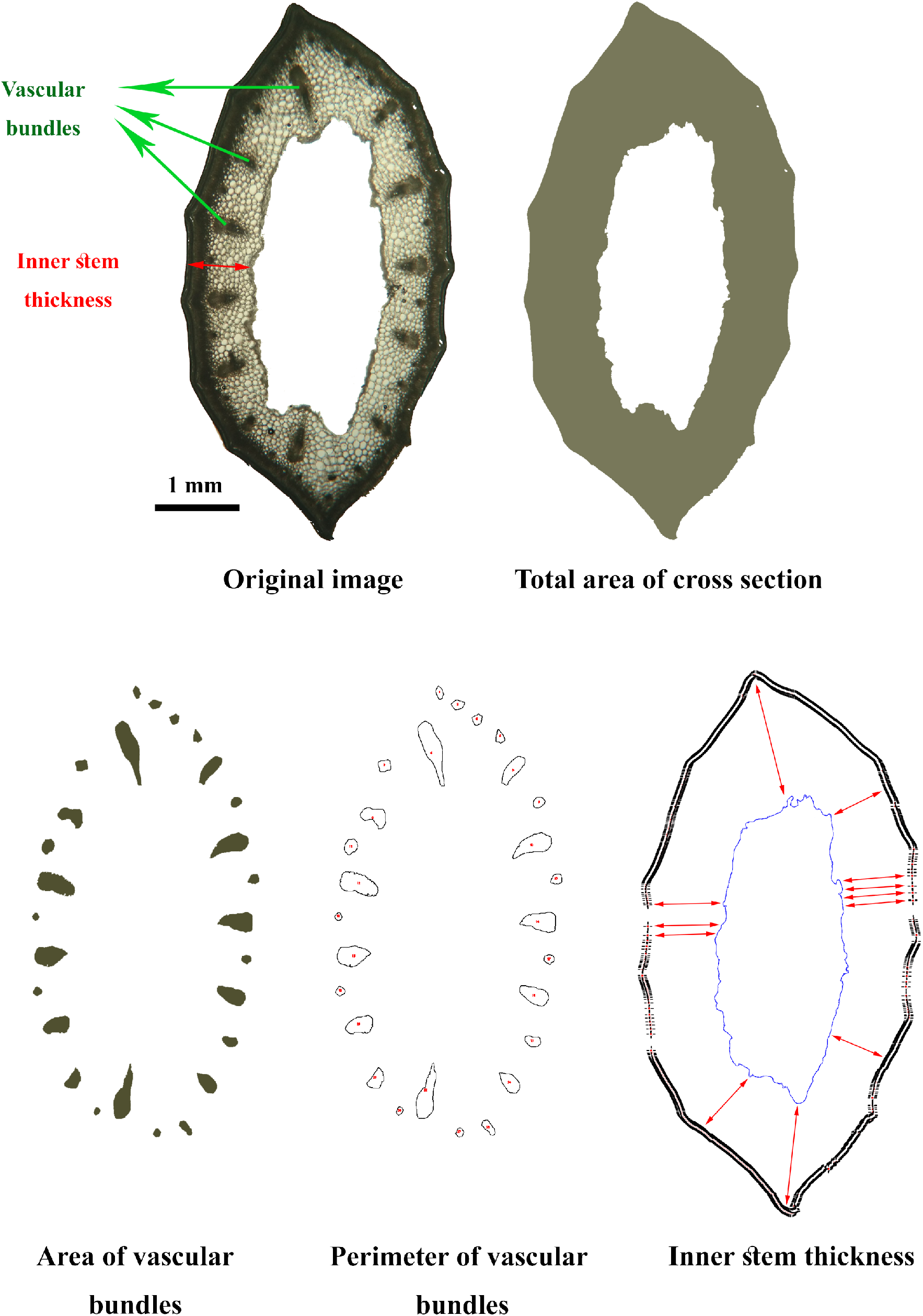
Measured indices related to the cross-sectional shape of narcissus flower stems.

For a precise definition of these indices, refer to the <u>ImageJ User Guide</u>. Generally, the Min Feret and Minor indices represent the width or smallest diameter of the object, whereas the Max Feret and Major indices indicate the length or largest diameter of the object.

## 2-5 Measurement of Biochemical Indices and Antioxidant Enzyme Activity

## 2-5-1 Mucilage Sugars in Stems

Immediately after harvest, the stems of cut flowers were separated and weighed with a digital scale accurate to two decimal places. The stems were longitudinally cut with a clean, sharp blade, and the soft tissue was scraped to remove all mucilage, which was collected in a clean container and weighed with the same scale. Mucilage extraction was performed for twelve stems from each genotype. The percentage of mucilage in the stem was calculated by dividing the mucilage weight by the stem weight. Then, 0.5 grams of mucilage was diluted with 1.5 ml of ultrapure water (Millipore) and mixed for two minutes. The measurement of mucilage sugars was performed in triplicate. The sugar content was measured via a high-performance liquid chromatography (HPLC) system with a refractive index (RI) detector. The diluted samples were filtered through a 0.45-micron filter and injected into the HPLC system with a 20-microliter volume. Sugar separation was performed at 65°C via a Eurokat pb-300 mm × 8 mm Knauer column with deionized water as the mobile phase and a flow rate of 1.8 millilitres per minute through a Smartline 1000 KNAUER pump. The concentrations of sucrose, glucose, and fructose were calculated via calibration curves of pure standards.

## 2-5-1 Malondialdehyde (MDA) and hydrogen peroxide (H_2_O_2_)

For the measurement of H_2_O_2_ and malondialdehyde (MDA) contents, the corona and petal tissues were carefully separated on Day 0 and Day 3, immediately frozen in liquid nitrogen, and stored at - 80°C. The extraction of H_2_O_2_ and MDA was performed with slight modifications to the method described by Zanganeh et al. (2019). A total of 0.1 grams of tissue was powdered in liquid nitrogen, and 2 mL of trichloroacetic acid (TCA) was added and homogenized. The homogenate was centrifuged at 10,000 rpm for 10 minutes, and the supernatant was used to measure the MDA and H_2_O_2_ levels. MDA was measured via the method of Hodges et al. (1999), and H_2_O_2_ was measured according to Alexieva et al. (2001).

## 2-5-2 Carotenoids, soluble proteins, and antioxidant enzyme activity

To measure total protein and antioxidant enzyme activities, fresh corona and petal tissues were quickly and carefully separated on days 0 and 3. The samples were immediately frozen in liquid nitrogen and stored at -80°C until enzyme assays were performed. Enzyme extraction was carried out with slight modifications to the method described by Sun et al. (2013). In this method, 0.5 grams of corona or petal tissue was powdered in liquid nitrogen, and 2 mL of cold 50 mM potassium phosphate buffer (pH 7) containing 1 mM EDTA and 1% (W/V) polyvinylpyrrolidone (PVP) was added and homogenized. The homogenate was centrifuged at 13,000 rpm for 10 minutes at 4°C, and the supernatant was used to determine the enzyme activity and total protein content.

The total protein content and antioxidant enzyme activities of peroxidase (POD), superoxide dismutase (SOD), and catalase (CAT) were measured spectrophotometrically. POD activity was measured via guaiacol, SOD activity was assessed via nitro blue tetrazolium (NBT), and CAT activity was determined via H_2_O_2_ (Zou et al., 2017). The total protein content was measured via the Bradford method (Bradford, 1976) with Coomassie Brilliant Blue.

Corona and petal tissues were carefully separated on the day of flower harvest, frozen in liquid nitrogen and stored at -80°C until carotenoids were measured. The total carotenoid content was measured via DMSO and spectrophotometry (Haghshenas et al., 2024).

## 2-6 Bacterial culture

To assess bacterial colony growth in the vase water, bacterial cultures were prepared. One milliliter of the vase solution was spread evenly on the surface of nutrient agar plates and incubated at 30°C for 48 hours (Rabiza-Świder et al., 2020a).

## 2-7 Data analysis

This study was conducted with a completely randomized design (CRD). The treatments included ten populations of Iranian narcissus. Analysis of variance (ANOVA) was performed via SPSS version 21; mean comparisons were carried out via Duncan’s multiple range test at the 5% significance level. Pearson correlation analysis was conducted via XLSTAT version 2016.02.28451, and a heatmap was created via the link http://www.heatmapper.ca/.

## 3 Results

## 3-1 Flower longevity

The longevity of flowers among the Iranian narcissus populations varied (Figures 2 and 3), as did the longevity of different floral tissues (Figure 3). White narcissus had the shortest floral longevity, with an average of 2.7 days, whereas the populations of Shahla from Khusf and Porpar from Khafr had the longest floral longevity, averaging 5.6 and 5.3 days, respectively. On average, the longevity of the corona was 6.6 days, and the longevity of the petals was 3.2 days. The shortest and longest average petal longevity were observed for white narcissus, with 2.2 days, and for the Porpar of Khafr and Khusf, with 4.8 and 4.6 days, respectively. The corona longevity varied among the populations, with the Shahla and Porpar populations having slightly longer than 7 days, while the white narcissus population had 3.8 days, and the Sadpar narcissus population had 4.5 days. Overall, the longevity of different floral tissues in narcissus varies (Figure 3), with petal senescence being the limiting factor for the longevity of the inflorescence.

**Fig. 2.**
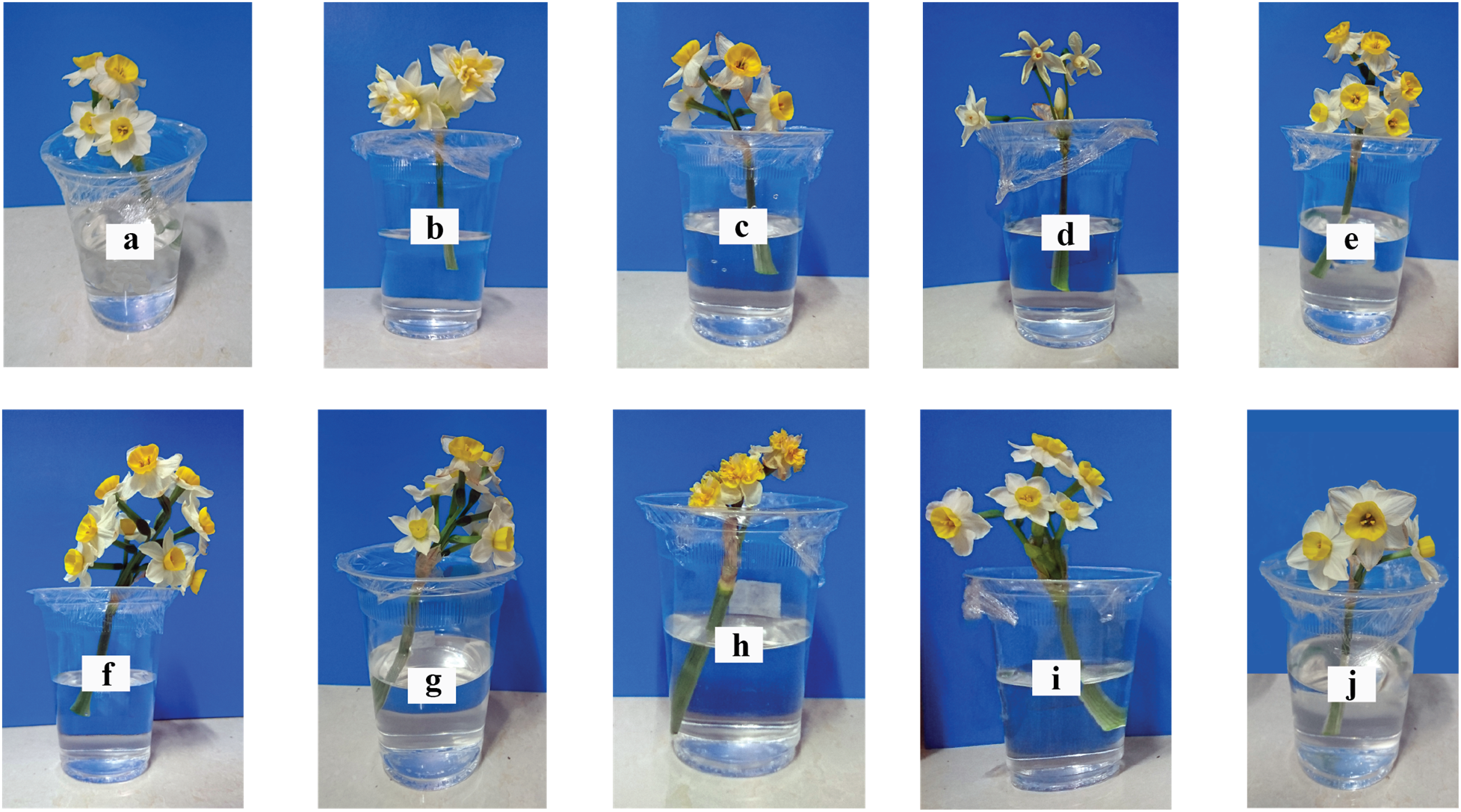
Iranian narcissus populations on the third day after harvest; a: Shahla Khafr, b: Porpar Khafr, c: Shahla Shiraz, d: White narcissus Shiraz (*N. papyraceus*), e: Shahla Abdanan, f: Shahla Kazerun, g: Shahla Behbahan, h: Sadpar Behbahan, i: Shahla Juybar, j: Shahla Khusf.

**Fig. 3.**
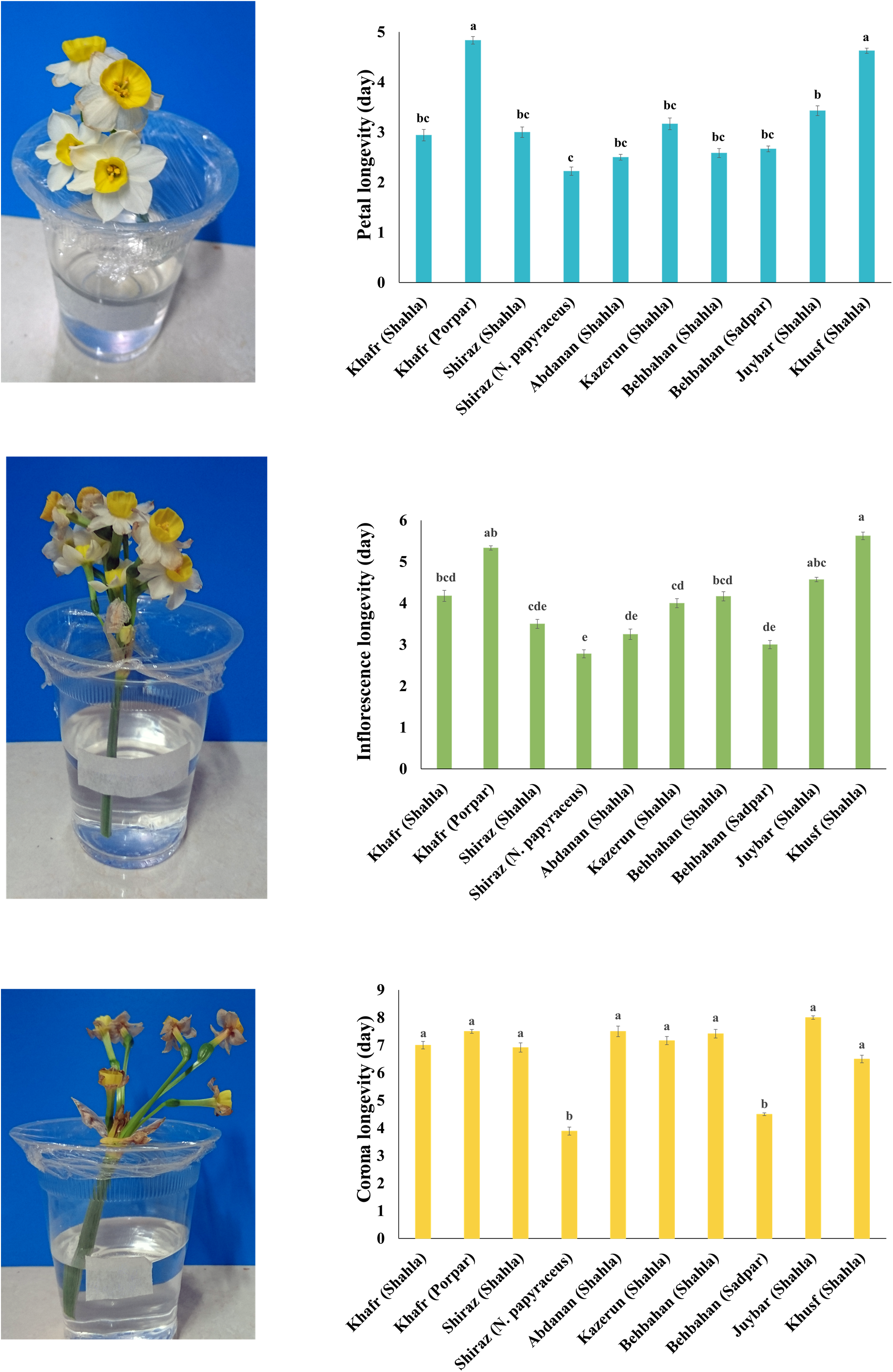
Longevity of inflorescence, coronas, and petals of Iranian narcissus populations.

## 3-2 Physiochemical Factors

The protein content in both the corona and petal tissues did not significantly differ; however, over time, the total protein content in both tissues significantly decreased (Figure 4). On the first day, the activity of the enzyme superoxide dismutase (SOD) was the same in both corona and petal tissues. By the third day, SOD activity significantly increased in the corona and decreased in the petals (Figure 4). From the start of the experiment, petal tissues presented significantly greater peroxidase activity than corona tissues did, and by the third day, peroxidase activity significantly increased in both tissues, with a greater increase observed in petals (Figure 4). There were significant differences in peroxidase activity among the populations of Iranian narcissus. *Narcissus papyraceus* had lower peroxidase activity than the other populations did, with a significant difference only observed in the Shahla Khafr population (data not shown). The electrical conductivity of both tissues was significantly greater on the third day than on the first day of the experiment. Three days after placing the inflorescences in the water, the H_2_O_2_ levels in both the corona and petal tissues significantly increased; on the third day, the H_2_O_2_ levels were greater in the petals than in the corona (Figure 4). There were significant differences in H_2_O_2_ levels among the Iranian narcissus populations, with *Narcissus papyraceus* showing the highest H_2_O_2_ levels, which significantly differed from those of the Shahla Khafr and Shahla Juybar populations (data not shown). The malondialdehyde (MDA) level was greater in petals than in coronas, and over three days, the MDA level in petals significantly increased, whereas the increase in the corona density was not significant. After three days, the relative water content (RWC) significantly decreased in both the corona and petal tissues. Although there were no significant differences in RWC among the narcissus populations, the white (*N. papyraceus*) population presented the lowest RWC (data not shown). In the narcissus populations, both the fresh and dry weights of the corona and petals significantly decreased after three days (Figure 4).

**Fig. 4.**
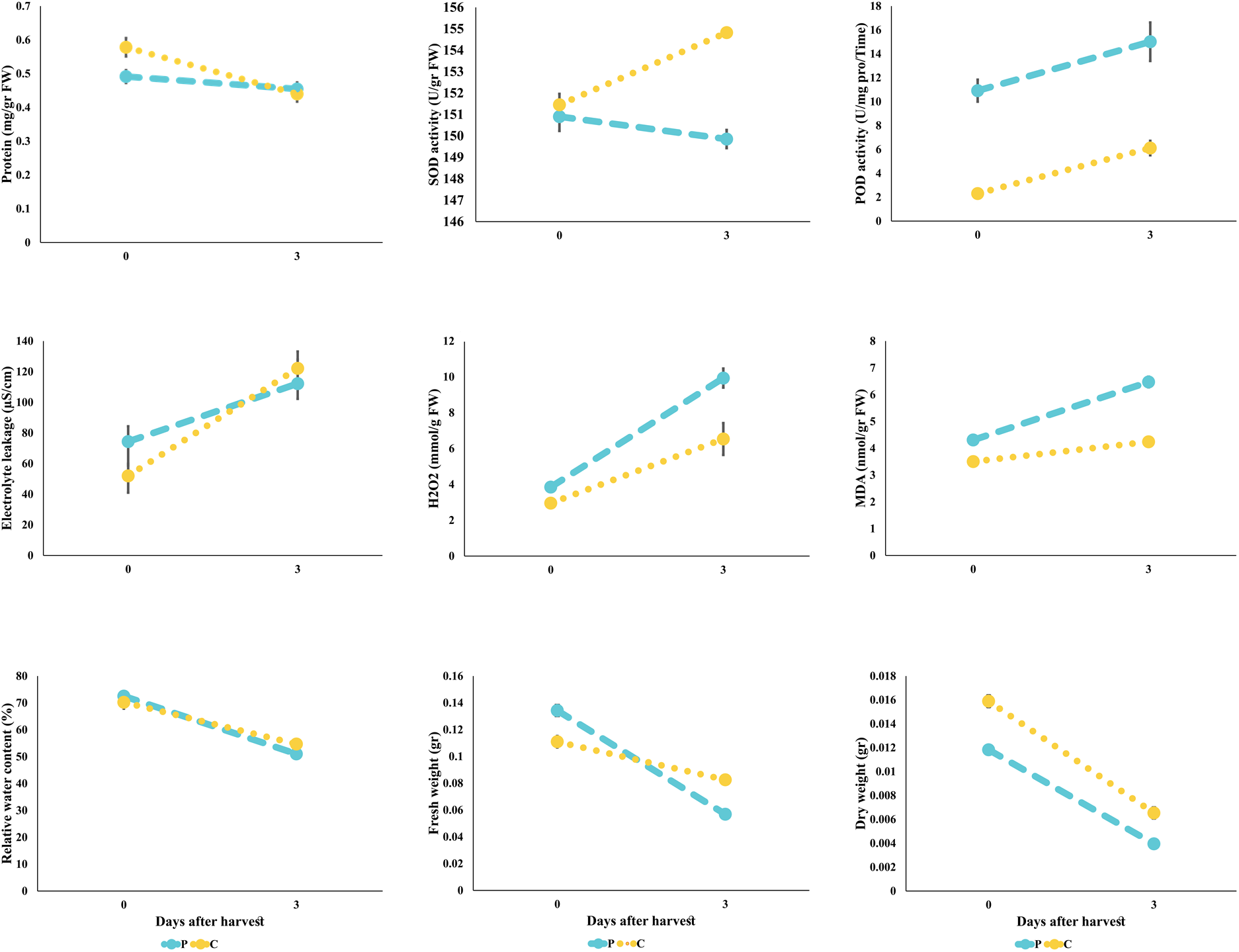
Changes in physicochemical factors in the corona and petal tissues of Iranian narcissus plants after three days of storage.

## 3-3 Vascular Bundles and Stems

Microscopic analysis of cross-sections of stems and vascular bundles revealed that the cross-sectional area of vascular bundles, cross-sectional area of the stem, stem perimeter (Figure 5), and stem solidity were significantly lower on the third day than on the first day (Figure 5). On the third day, the Max Feret (largest diameter), Min Feret (smallest diameter), Minor, and Major of the stems were significantly lower than those on the first day (Figure 5).

**Fig. 5.**
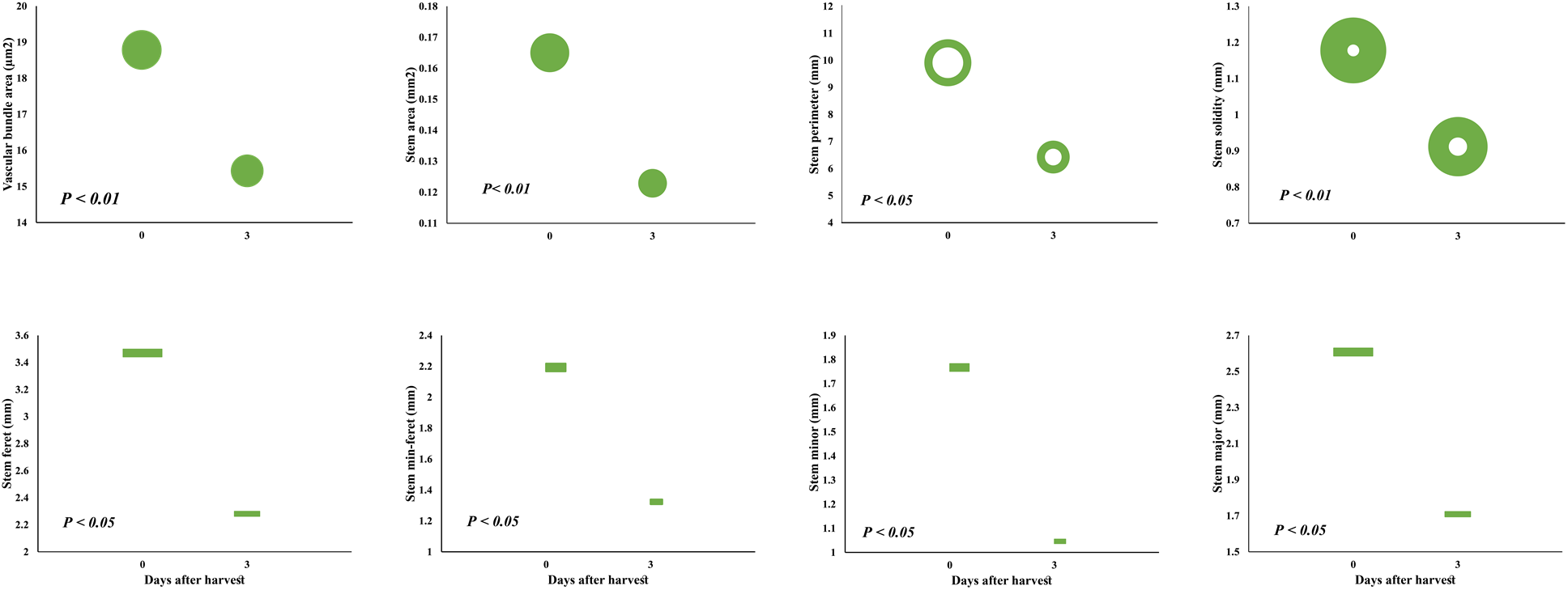
Quantitative changes in the stem and vascular bundles of Iranian narcissus after three days of storage in vase.

No vascular blockage was observed in the stems of narcissus on the third day (Figure 6). In the bacterial cultures, there were no significant differences in the number of bacteria grown in the floral preservative solutions among the different genotypes. However, there was a significant difference in the number of bacterial colonies growing in the vase solutions of the Shahla from the Kazerun, Khusf, and Behbahan genotypes compared with the control (not shown). The amount of sucrose in the stem mucilage had a small but significant correlation with bacterial growth (R = 0.373), whereas fructose and glucose did not correlate with bacterial growth.

**Fig. 6.**
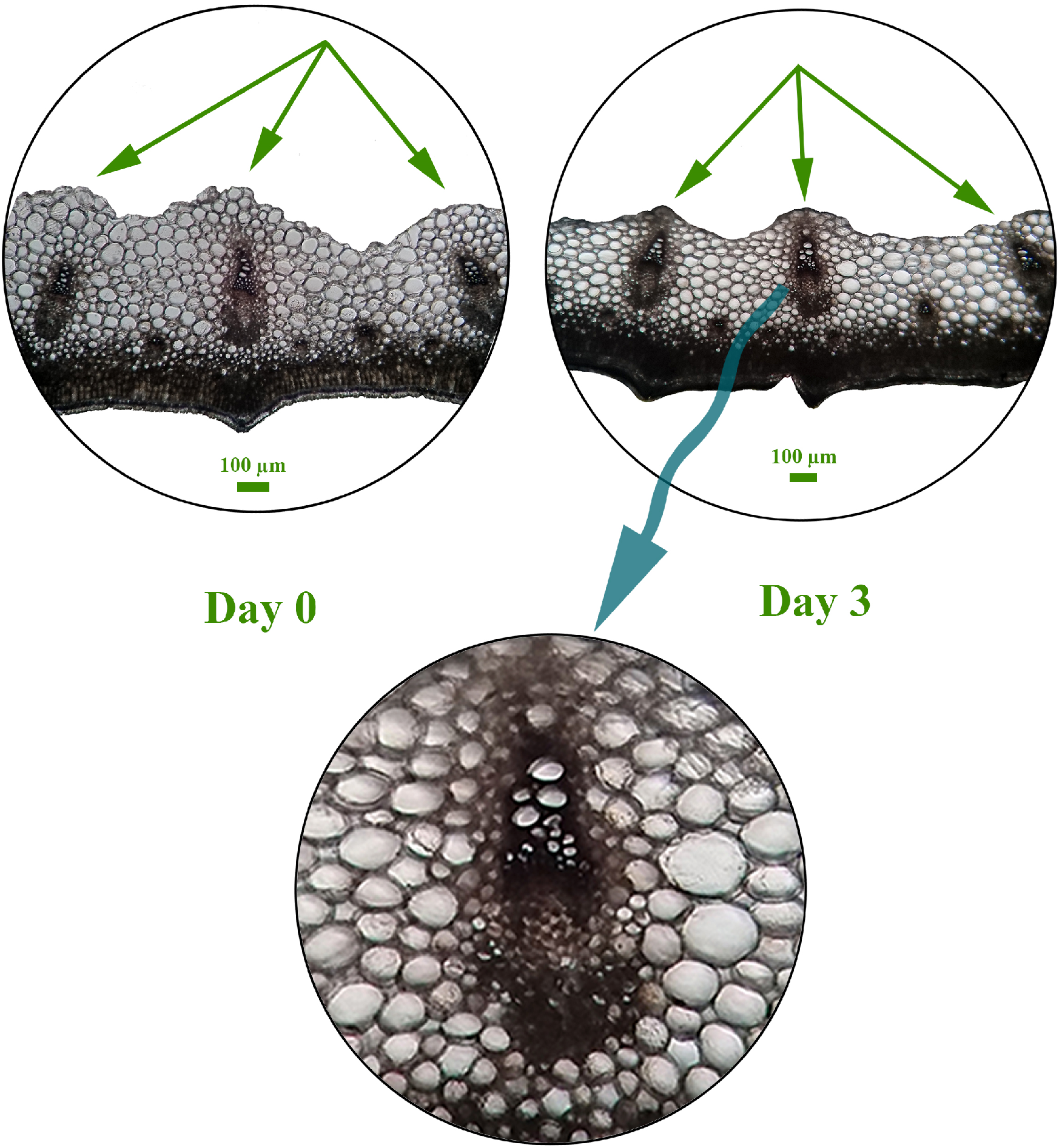
Microscopy images of vascular bundles in the stem of Iranian narcissus in freshly harvested stems and after three days of storage in vase. Microscopy images were taken with a polarized light microscope and a mobile phone camera. The lack of vascular blockage on the third day is evident in the lower image. The green arrows indicate the collapse of stem tissue cells near the central cavity of the stem.

## 3-4 Relationships among flower longevity, vascular traits, and physiochemical factors

The cross-sectional area of vascular bundles, the fresh weight of petals, and the amount of H₂O₂ in the corona affected flower longevity. Specifically, flower longevity was positively correlated with the cross-sectional area of vascular bundles on the third day and the fresh weight of petals on the first day, whereas it was negatively correlated with H₂O₂ levels in the corona on both the first and third days (Figure 7). The longevity of the corona was positively correlated with the number of vascular bundles, the cross-sectional area of the vascular bundles on both the first and third days, and the RWC on the third day (Figure 7). The number of vascular bundles was positively correlated with the fresh weight of the petals and the corona (Figure 7). Additionally, the number of vascular bundles was highly significantly positively correlated with relative water uptake (RWU; Figure 8).

**Fig. 7.**
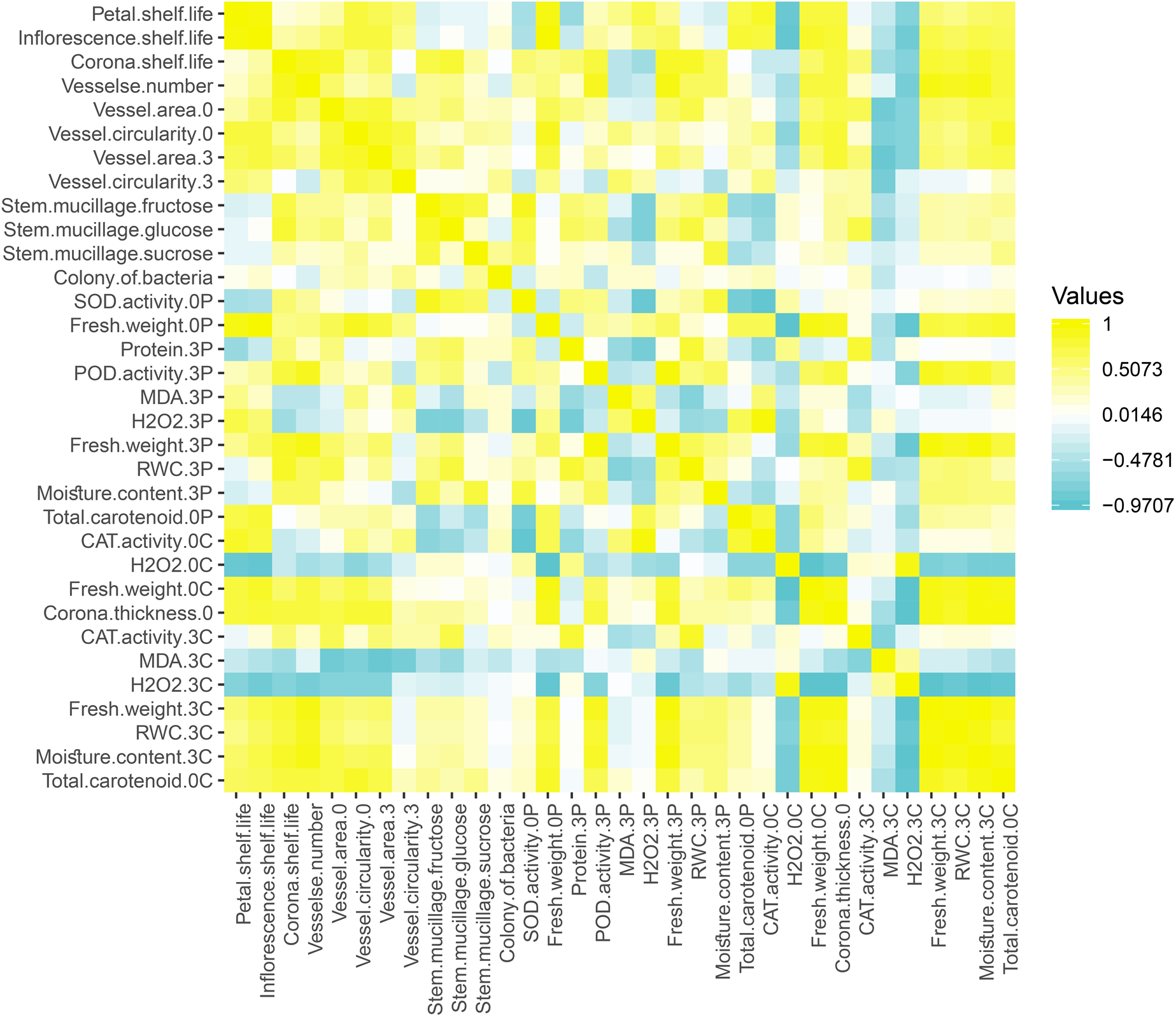
Heatmap of flower longevity, vascular traits, and physiochemical factors in Iranian narcissus.

**Fig. 8.**
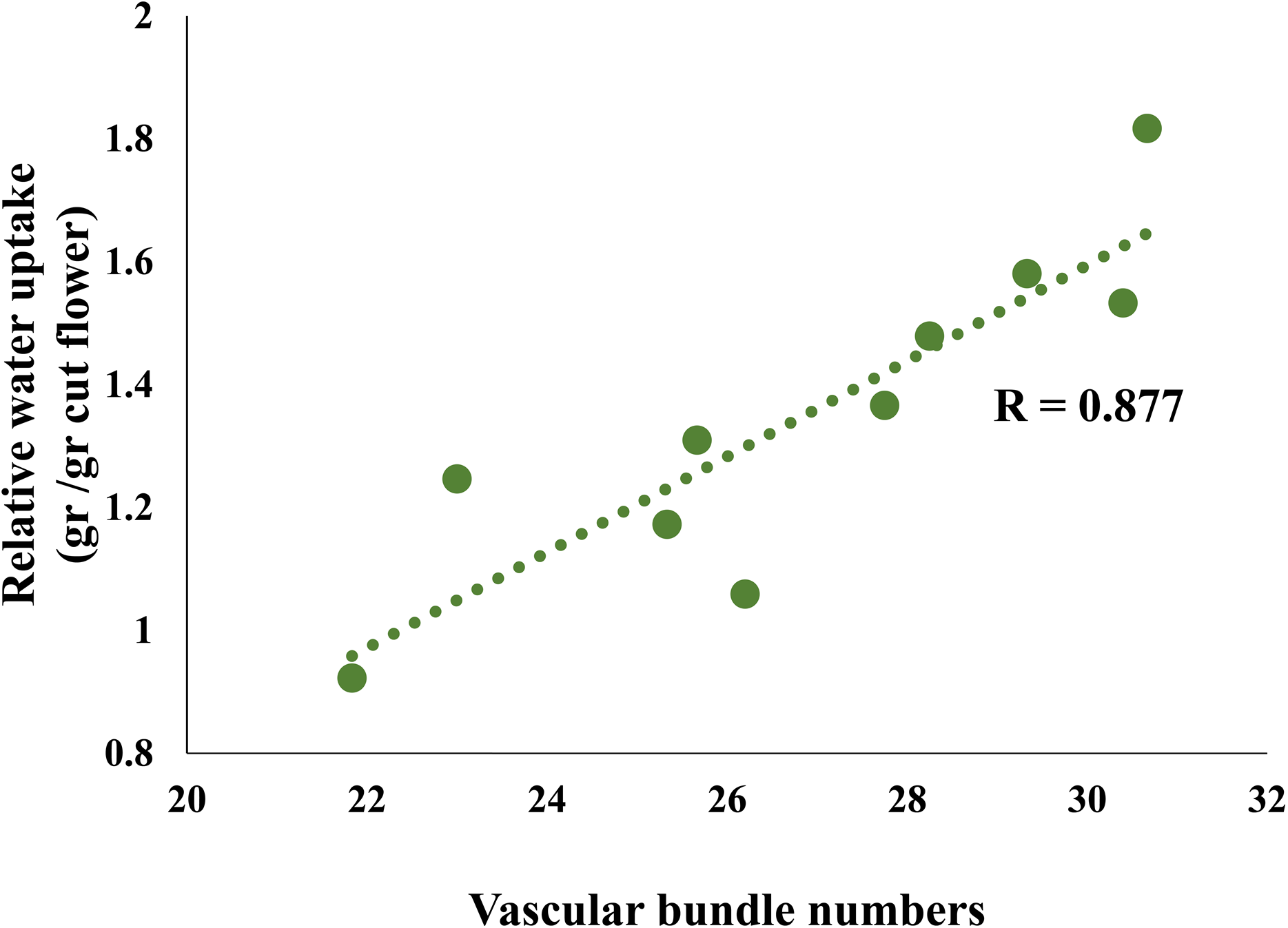
Significant correlation between relative water uptake and the number of vascular bundles in Iranian narcissus populations.

On the third day, the protein content in the petals was significantly positively correlated with the RWC, and POD enzyme activity was significantly positively correlated with fresh weight. The correlation between the malondialdehyde (MDA) content and the RWC of the petals on the third day was significantly negative. On both the first and third days, H₂O₂ levels were significantly negatively correlated with the fresh weight of the corona (Figure 7). The carotenoid content of the corona was significantly negatively correlated with H₂O₂ levels and significantly positively correlated with the percentage of moisture and fresh weight of the corona on the third day. Additionally, the fresh weight of the corona significantly positively correlated with the corona thickness on the first day.

## 4 Discussion

## 4-1 Flower longevity in Iranian narcissus populations

Flower longevity in narcissus was genotype dependent. The average longevity of the Shahla narcissus populations without any treatment or recutting was 4.18 days; Porpar narcissus had 24% longer longevity, while Sadpar narcissus had 32% shorter longevity, and White narcissus (*N. papyraceus*) had 40% shorter longevity, compared with Shahla narcissus. The white narcissus presented the highest levels of H₂O₂ and the lowest RWC, peroxidase activity, and carotenoid content in the corona tissue, which was correlated with the shorter flower longevity of this population. The genetic characteristics of plants are significant factors in cut flower longevity (Vijayakumar et al., 2021). Previous studies reported average narcissus flower longevity ranging from 4 to 6 days (Rabiza-Świder et al., 2020c). Barfi et al. (2021) reported varying flower vase life among Iranian narcissus populations in field conditions in Ahvaz, attributing this variation to diverse climatic conditions in Iran and high mutation rates in these populations (Barfi et al., 2021). Their study revealed that the Abdanan population had the longest vase life, whereas in the current study, the Khusf and Khafr populations had the longest longevity. This discrepancy may be due to different environmental conditions affecting postharvest flower life and variations in the genotypes examined from those regions (Barfi et al., 2021).

Research on chrysanthemum has shown that the varying vase life among different cultivars of cut chrysanthemum is due to differences in oxidative status and physiological responses to postharvest conditions (Fanourakis et al., 2022). Water relations in flower vase life plays a lesser role than does oxidative stress in determining flower longevity (Fanourakis et al., 2022). Studies on different rose cultivars have reported morphological differences, such as differences in stomatal number and distribution, and physiological behaviors, such as the ability of cultivars to reduce transpiration by closing their stomata (Wolteringa and Paillarta, 2018; Fanourakis, 2020).

Rabiza-Świder et al. (2020a) reported that the longevity of 14 peony cultivars was affected differently by postharvest treatments involving 8-hydroxyquinoline citrate (8-HQC), nanosilver, and sucrose and that postharvest treatments should be tailored to each cultivar separately (Rabiza-Świder et al., 2020a). The postharvest longevity of cut flowers depends on the genetic and heritable traits of the plant and is influenced by environmental factors (Verdonk et al., 2023; Rabiza-Świder et al., 2020b; Vijayakumar et al., 2021). In the present study, flower longevity was positively correlated with the cross-sectional area of vascular bundles, and the number of vascular bundles was positively correlated with the fresh weight of the corona and petals and the relative water uptake (RWU). It appears that genotypes with a greater number of vascular bundles, which facilitate greater water uptake and transport to flower tissues, experienced delayed stress and free radical production.

## 3-4-2 Senescence process and physiological factors in cut narcissus flowers

More than 25% of postharvest changes in flowers are related to signs of aging (Rabiza-Świder et al., 2020b). Aging is a programmed process that does not occur simultaneously in all floral organs. Owing to their specific biological functions, petals are the first tissues to show signs of aging, such as wilting and color change (Hong-mei Fan et al., 2015; Sun et al., 2021). Cell and tissue death within a flower occurs at multiple levels (Rabiza-Świder et al., 2020c). In this study, the first visible signs of aging appeared as browning of petal margins and ended with drying and color changes in the corona; overall, the corona lasted twice as long as the petals did. Rabiza-Świder et al. (2020c) reported that the average longevity of petals for various Dutch narcissus cultivars was 4 to 5 days, whereas the average longevity of petals in coronas ranged from 6 to 7 days (Rabiza-Świder et al., 2020c). Previous research on Iranian narcissus populations (Haghshenas et al., 2024) measured the thickness of the corona and petal tissues; petal thickness was 48% less than that of the corona. Thus, the corona has a lower surface-to-volume ratio and greater moisture retention capacity compared with the petals. One reason for the greater longevity of the corona compared with that of the petals may be the lower rate of evaporation from the corona tissue, such that by the third day, the fresh weight loss of the petals was 92% greater than that of the corona. Rabiza-Świder et al. (2020c) reported that the RWC in the corona tissue of Dutch narcissus was greater at harvest than in that of petals and that the turgidity of epidermal cells in the corona and petals of Dutch narcissus disappeared a few days after placement in flower preservative solution, leading to cell collapse and signs of wilting. The resistance of corona epidermal cells to these changes is greater than that of petals, which is why the corona has longer longevity (Rabiza-Świder et al., 2020c). The ability to uptake water from the preservative solution, water loss through transpiration, and the ability of the cut flower stem to retain water are three key factors affecting the water balance of cut flowers. When the balance between water uptake by the stem and water loss through transpiration is disrupted, wilting, a visible sign of aging and the end of flower vase life, occurs (Manzoor et al., 2024). Additionally, Rabiza-Świder et al. (2020c) reported that the sugar content in the ovary and corona tissue increased after placement in flower preservative solution, whereas the sugar content in the petals did not change (Rabiza-Świder et al., 2020c), which may also explain the greater longevity of the corona than that of the petals.

Compared with the corona, the petals had higher levels of H₂O₂ and MDA; on the third day, the petals had 41% more H₂O₂ and MDA than did the corona. Generally, the correlation between H₂O₂ and the fresh weight of tissues indicates that H₂O₂ production in corona and petal tissues could be one reason for the reduced fresh weight. Oxidative stress leads to imbalance between the accumulation of reactive oxygen species (ROS), such as hydrogen peroxide (H₂O₂), and that of antioxidants (Fanourakis et al., 2022). Membrane lipid peroxidation and increased membrane permeability are associated with aging processes. Increased membrane permeability leads to cell sap leakage and eventually to cell death. Lipid peroxidation intermediate products, free radicals, and the final product, malondialdehyde, accumulate in tissues during aging. The levels of malondialdehyde and free radicals such as H₂O₂ are important indicators of tissue damage during aging (Zhao et al., 2018; Rabiza-Świder et al., 2020b; Gururani et al., 2023), and the levels of these compounds often increase in cut flowers after detachment from the mother plant (Rabiza-Świder et al., 2020b). In some cut flowers, such as roses, peonies, and irises, these indicators are considered signs of flower aging (Rabiza-Świder et al., 2020a). In cut orchids, one indicator of aging is increased H₂O₂ in pollinated flowers (Rabiza-Świder et al., 2020b). According to Fanourakis et al. (2022), H₂O₂ accumulation increases lipid membrane oxidation during the storage of cut chrysanthemums. Increased lipid oxidation results in increased production of malondialdehyde (Fanourakis et al., 2022). In roses, H₂O₂ production increases until the eighth day and then decreases, and this increase in H₂O₂ is associated with higher malondialdehyde levels (Mazrou et al., 2022). In the daylily, gladiolus, and snapdragon, H₂O₂ begins to increase before the full development of the flower and reaches its peak when the flower is fully open. In snapdragon, younger florets at the top of the inflorescence had significantly less H₂O₂ than older florets at the bottom (Rabiza-Świder et al., 2020b). In the present study, visible signs of aging in narcissus flowers began with petals, and the higher levels of H₂O₂ and MDA in the petals appear to contribute to their earlier aging. On the basis of our observations, in narcissus, petals wilt first even when attached to the mother plant, and the onset of visible signs of aging coincides with petal aging, which is characteristic of this plant. Morphologically, petals and the corona are similar, but their origin and timing of differentiation differ. The corona’s differentiation is independent of the usual four floral whorls (sepals, petals, stamens, and pistils) and occurs after the differentiation of all flower parts; structurally (Ma et al., 2023) and in terms of ABC gene expression involved in floral tissue differentiation (Waters et al., 2013), it resembles stamens. Petals play a biological role in attracting pollinators for the reproductive process and start aging after pollination, turning into a nutritional source for reproductive organs (Sun et al., 2021). The corona, which resembles the reproductive structure of stamens, has a longer lifespan than petals do and plays a critical role in attracting pollinators because of its color and shape; thus, it naturally has longer longevity than petals do. This feature was also observed under postharvest conditions. Therefore, it seems that the increase in free radicals in the petals occurs earlier than that in the corona, even under non-postharvest conditions.

Plants protect their cells from free radicals via an antioxidant defense system. The activity of protective enzymes increases during aging (Rabiza-Świder et al., 2020b). In this study, the superoxide dismutase activity and dry weight of the petals were lower than those of the corona. The antioxidant enzyme activities in the corona and petals were not the same; over the initial three days of the experiment, peroxidase activity increased in the petals, whereas superoxide dismutase activity increased in the corona. Mazrou et al. (2022) reported an increase in antioxidant enzyme activity until the sixth day in roses, followed by a decrease (Mazrou et al., 2022). Rabiza-Świder et al. (2020b) reported that, three days after placement in a flower preservative solution, snapdragon flowers presented increased catalase activity in younger florets at the top of the inflorescence and decreased activity in older florets at the bottom (Rabiza-Świder et al., 2020b). As stress conditions begin, increasing levels of ROS activate antioxidant enzymes to protect flowers from ROS damage, but with increasing stress conditions, excessive ROS accumulation inhibits the effectiveness of the antioxidant defense system (Rabiza-Świder et al., 2020b; Zhao et al., 2018). Another crucial component of the antioxidant defense system is flower pigments such as carotenoids (Cavaiuolo et al., 2013), which play an antioxidant role in plants and animals (Parklak et al., 2023; Cavaiuolo et al., 2013; Cavallaro et al., 2023), and their levels vary among species and even cultivars (Cavaiuolo et al., 2013). In this study, White narcissus had the lowest carotenoid content in its corona (Haghshenas et al., 2024) and also had the shortest flower longevity among the narcissus populations examined. Additionally, the yellow coronas of the other genotypes contained significantly greater levels of carotenoids compared with the white petals. These findings suggest that the greater longevity of the corona may also be attributed to the role of carotenoids in scavenging free radicals.

Three days after harvest, protein levels decreased in both the corona and petal tissues of narcissus flowers. Typically, proteins in cut flowers begin to degrade after detachment from the mother plant, and their levels decrease with wilting (Rabiza-Świder et al., 2019). Studies have shown that even in unharvested flowers, soluble protein levels decrease during natural aging. This reduction is due to decreased protein synthesis and increased degradation by various proteases. Protein levels decrease in flowers attached to the mother plant before opening, but in harvested cut flowers, protein loss occurs after flower opening (Gul et al., 2020).

## 4-3 Water stress and vascular occlusion in cut narcissus flowers

Because cut flowers are separated from their mother plant, they experience water stress. During the postharvest period, the xylem vessels at the end of the stem begin to become blocked due to bacterial growth (Rabiza-Świder et al., 2020b, 2020c) and physiological responses of the stem to wounding, such as the deposition of gums, mucilage (Verdonk et al., 2023), extracellular polysaccharides, cellular degradation products (Rabiza-Świder et al., 2020c), and the formation of tyloses (Verdonk et al., 2023). This blockage can reduce water uptake and transport, leading to premature wilting of flowers (Rabiza-Świder et al., 2020b). Skutnik et al. (2020) reported that after 12 weeks of storage, no xylem blockage was observed in the stems of cut peony flowers (Skutnik et al., 2020). In the present study, no vascular occlusion was observed in narcissus stems three days after being placed in a floral vase. However, cell collapse at the cut site led to a reduction in the thickness of the parenchyma tissue and subsequently reduced the perimeter, area and solidity of the stem at the cut site. Structural changes associated with programmed cell death during the aging process, such as the collapse of mesophyll and epidermal cells and a reduction in nucleus size, have been observed under a microscope in various flowers, such as iris, alstroemeria, gypsophila, jasmine, and clematis (Rabiza-Świder et al., 2020b). In gypsophila, mesophyll cells begin to collapse even before the first signs of aging are visible (Rabiza-Świder et al., 2020b). Essentially, the aging process in narcissus flowers begins at the cut site and continues as the stem tissues are immersed in the vase solution. Cutting the stem causes oxidative damage and excessive production of reactive oxygen species (ROS), which attack cellular proteins, nucleic acids, and membrane lipids, leading to membrane degradation (Mazrou et al., 2022). Over time, these stresses result in the collapse of the cells at the cut end of the stem.

In this study, a positive correlation was detected between the number of bacteria growing in the vase solution and the sucrose content in the stem mucilage. These findings indicate that the sucrose released from the cut stem into the vase solution promoted bacterial colony growth in the solution. Additionally, there was a significant difference in the number of bacterial colonies among the different narcissus genotypes from the kazerun, Khusf, and Behbahan populations compared with the control; however, no vascular occlusion was observed in the stems of the narcissus flowers studied. The mucilage of the Dutch narcissus stem contains sugar and alkaloid compounds, which, when placed together with other flowers in a vase, have varying effects on their longevity. (Van Doorn, 1998; Van Doorn et al., 2004). Van Doorn (1998) studied the effect of mucilage secreted from the stem of Dutch narcissus on the longevity of rose and tulip flowers. He reported that mucilage secreted from narcissus stems, due to increased sugar content in the preservative solution and thus increased bacterial growth, led to vascular occlusion in rose stems and decreased flower longevity. In tulips, although no vascular occlusion was observed due to bacterial growth, the alkaloid compounds secreted from narcissus stems had a toxic effect that reduced the longevity of tulips (Van Doorn, 1998). In the present study, although the sucrose in the stem mucilage that was released into the distilled water of the vase increased the number of bacterial colonies, these colonies did not cause vascular occlusion. Both tulips and narcissus are herbaceous bulbous plants and may have developed similar mechanisms to prevent vascular occlusion. In floral preservative solutions, sucrose is commonly used to compensate for the carbohydrates consumed during respiration and to regulate the osmotic pressure of the solution, but sucrose use in preservative solutions can lead to increased bacterial colony counts (Rabiza-Świder et al., 2020a and 2020c). Bacteria in the preservative solution likely originate from the stem surface and utilize carbohydrates leaching from wounded surfaces of the stem and also from removed leaves and thorns (Verdonk et al., 2023). Rabiza-Świder et al. (2020c) reported that after four to five days of holding Dutch narcissus flowers in preservative solutions, no vascular occlusion was observed (Rabiza-Świder et al., 2020c). Vascular occlusion depends on the flower species and variety (Van Doorn, 1998; Van Doorn et al., 2004). Wolteringa and Paillarta (2018) reported that regardless of the presence of bacteria in preservative solution, the storage of flowers leads to a reduced water content in the tissues and consequently decreases flower longevity, and bacterial colony growth alone does not increase the resistance of xylem vessels to water uptake (Wolteringa & Paillarta, 2018). In addition to the different responses of species to vascular occlusion caused by bacterial growth, the effects of alkaloid toxicity are not uniform across all cut flowers. Van Doorn et al. (2004) investigated the effect of narcissus stem mucilage on the longevity of cut iris flowers by placing Dutch narcissus flowers in a preservative solution with irises and treating the solution with mucilage and a low concentration (100 mg L^-1^) of narciclasine, the predominant alkaloid in Dutch narcissus. They reported that the mucilage secreted from the narcissus stem delayed the onset of senescence symptoms in iris flowers by preventing the synthesis of proteases, attributing this effect to the narciclasine present in the mucilage (Van Doorn et al., 2004). Tarakemeh et al. (2019) reported no narciclasine among the ten alkaloids identified in five Iranian narcissus varieties (*N. tazetta* and *N. papyraceous*; Tarakemeh et al., 2019). The exact reasons for the lack of vascular occlusion by bacteria or physiological blockage in the narcissus flowers studied in this research are unclear. Possible explanations include the following: 1) whether the bacterial count in the preservative solution was insufficient for vascular blockage; 2) whether other compounds in the stem mucilage can prevent vascular occlusion; and 3) whether, given the monocot and herbaceous nature of narcissus, there is a different physiological response than that of dicot plants regarding wounds and the formation of tyloses, calluses, and substances that cause vascular occlusion.

The sucrose content in the stem mucilage was also significantly positively correlated with the moisture percentage in the petal tissue on the third day. Genotypes whose stem mucilage had relatively high sucrose levels presented relatively high moisture percentages in their petals after three days. As mentioned, one of the roles of sugars is osmotic regulation. Sugars in the preservative solution are transferred to the petals, aiding in carbohydrate accumulation, primarily as reducing sugars. These sugars act as osmolytes, increasing water flow into petals (Rabiza-Świder et al., 2020a and 2020c). Sucrose in stem mucilage also helps water absorption by regulating osmotic pressure and maintaining tissue hydration. Although the sugar content in the tissues does not always directly correlate with flower longevity (Rabiza-Świder et al., 2019), in bulbous plants, the application of sugars in the preservative solution does not increase flower longevity because sugars are consumed by other parts of the flower in addition to the petals. In narcissus flowers, where petals face energy shortages, sugars in the preservative solution are absorbed by highly metabolically active ovaries (Rabiza-Świder et al., 2020c).

In this study, the examination of structural changes in the stems of cut narcissus flowers during the postharvest period was performed via sectional imaging of microscopic samples without slide preparation, and the results were analyzed via ImageJ software. To the best of our knowledge, structural changes in the stem and vascular bundles of cut flowers, especially narcissus, during the postharvest period have not been previously reported. The use of phenotypic approaches and phenotyping methods can significantly aid in understanding the morphophysiological changes in plant organs during the postharvest period.

## 5 Conclusion

The longevity of Iranian narcissus cut flowers is influenced by the genotype and various morphological, physiological, and biochemical characteristics of the flowers. Overall, the postharvest life of narcissus flowers depends on the level of free radicals produced in the tissues; the activity of antioxidant defense systems, such as antioxidant enzyme activity and carotenoid content; the number of vascular bundles; and the sucrose content in the stem mucilage. This study examined the changes in stem tissue and vascular bundles during the postharvest period in cut narcissus flowers via microscopic imaging of stem cross-sections. Over time, the cells at the stem ends collapsed; it appears that the wound at the stem end, followed by exposure of the stem-end cells to the waterlogged environment of the preservative solution, created stress conditions at the stem ends, leading to cell death and collapse, which likely affected water uptake. Such changes in cut flowers, especially narcissus, have not been previously reported. The use of phenotyping methods and examination of the phenotypic characteristics of plant tissues can significantly aid in understanding the morphophysiological changes in plant organs during the postharvest period. In the studied narcissus cut flowers, no vascular occlusion was observed. The reason for the lack of vascular occlusion is not clearly understood—whether the compounds present in the stem mucilage prevent vascular occlusion or if the monocot nature of the plant results in different physiological responses to wounds than those of dicot plants. Additionally, the longevity of the petal and corona tissues of narcissus flowers is not the same. Aging signs appeared later in the corona tissues because of the following features: 1. The greater the thickness of the corona is, which reduces water loss compared with that of the petals; 2. Low levels of free radicals such as H_2_O_2_, 3. Better membrane integrity and lower malondialdehyde production; 4. Compared with petal tissue, it has a more robust antioxidant defense system with different antioxidant enzyme activities and higher carotenoid contents. By understanding the internal morphophysiological and biochemical changes in the inflorescence, more effective treatments for extending the longevity of cut flowers can be selected. Additionally, in breeding programs, considering the features that affect inflorescence longevity and selecting genotypes with longer-lasting flowers can help reduce postharvest losses and increase the profitability of the cut flower industry.

## Notes

### Competing Interest Statement

The authors have declared no competing interest.

## References

Alexieva, V., Sergiev, I., Mapelli, S., & Karanov, E. (2001). The effect of drought and ultraviolet radiation on growth and stress markers in pea and wheat. Plant, Cell & Environment, 24(12), 1337–1344. https://onlinelibrary.wiley.com/doi/epdf/10.1046/j.1365-3040.2001.00778.x.

Armitage, A. M. (1993). Specialty cut flowers. The production of annuals, perennials, bulbs and woody plants for fresh and dried cut flowers (pp. 384-pp).

Barfi, F., Salmi, M. S., & Zare, A. (2021). Investigation of morphological and biochemical traits charactristics related to vase life in population narcissus (*Narcissus tazetta* L.) in Khuzestan climate Iran. Iranian Journal of Rangelands and Forests Plant Breeding and Genetic Research, 29 (2), 282–296. (In farsi). https://doi.or/10.22092/IJRFPBGR.2022.357612.1407.

Bradford, M. M. (1976). A rapid and sensitive method for the quantitation of microgram quantities of protein utilizing the principle of protein-dye binding. Analytical Biochemistry, 72(1-2), 248–254. 10.1016/0003-2697(76)90527-3.

Cavaiuolo, M., Cocetta, G., & Ferrante, A. (2013). The antioxidants changes in ornamental flowers during development and senescence. Antioxidants, 2(3), 132–155. 10.3390/antiox2030132.

Da Silva, J. A. T. (2015). Ornamental cut flowers: physiology in practice. Floriculture & Ornamental Biotechnology, 124–140.

Darras, A. (2021). Overview of the dynamic role of specialty cut flowers in the international cut flower market. Horticulturae, 7(3), 51. 10.3390/horticulturae7030051.

Doorn, W. V. (1998). Effects of daffodil flowers on the water relations and vase life of roses and tulips. https://www.cabidigitallibrary.org/doi/full/10.5555/19980303976.

Fan, H. M., Li, T., Sun, X., Sun, X. Z., & Zheng, C. S. (2015). Effects of humic acid derived from sediments on the postharvest vase life extension in cut chrysanthemum flowers. Postharvest Biology and Technology, 101, 82–87. 10.1016/j.postharvbio.2014.09.019.

Fanourakis, D., Bouranis, D., Tsaniklidis, G., Rezaei Nejad, A., Ottosen, C. O., & Woltering, E. J. (2020). Genotypic and phenotypic differences in fresh weight partitioning of cut rose stems: Implications for water loss. Acta Physiologiae Plantarum, 42, 1–10. 10.1007/s11738020-03044-w.

Fanourakis, D., Papadopoulou, E., Valla, A., Tzanakakis, V. A., & Nektarios, P. A. (2021). Partitioning of transpiration to cut flower organs and its mediating role on vase life response to dry handling: A case study in chrysanthemum. Postharvest Biology and Technology, 181, 111636.

Fanourakis, D., Papadakis, V. M., Psyllakis, E., Tzanakakis, V. A., & Nektarios, P. A. (2022). The role of water relations and oxidative stress in the vase life response to prolonged storage: A case study in chrysanthemum. Agriculture, 12(2), 185. 10.3390/agriculture12020185.

Gul, F., Tahir, I., & Shahri, W. (2020). Flower senescence and some postharvest considerations of *Amaryllis belladonna* cut scapes. Plant Physiology Reports, 25, 315–324. 10.1007/s40502-020-00506-8.

Gururani, M. A., Atteya, A. K., Elhakem, A., El-Sheshtawy, A. N. A., & El-Serafy, R. S. (2023). Essential oils prolonged the cut carnation longevity by limiting the xylem blockage and enhancing the physiological and biochemical levels. Plos One, 18(3), e0281717. 10.1371/journal.pone.0281717.

Haghshenas, A., Jowkar, A., Chehrazi, M., Moghadam, A., & Karami, A. (2024). Fragrance and color production from corona and perianth of Iranian narcissus (*Narcissus tazetta* L.). Industrial Crops and Products, 212, 118368. 10.1016/j.indcrop.2024.118368.

Hodges, D. M., DeLong, J. M., Forney, C. F., & Prange, R. K. (1999). Improving the thiobarbituric acid-reactive-substances assay for estimating lipid peroxidation in plant tissues containing anthocyanin and other interfering compounds. Planta, 207, 604–611. https://link.springer.com/article/10.1007/s004250050524.

Kondo, M., Nakajima, T., Shibuya, K., & Ichimura, K. (2020). Comparison of petal senescence between cut and intact carnation flowers using potted plants. Scientia Horticulturae, 272, 109564. 10.1016/j.scienta.2020.109564.

Lear, B., Casey, M., Stead, A. D., & Rogers, H. J. (2022). Peduncle necking in *Rosa hybrida* induces stress-related transcription factors, upregulates galactose metabolism, and downregulates phenylpropanoid biosynthesis genes. Frontiers in Plant Science, 13, 874590. 10.3389/fpls.2022.874590.

Ma, Y., Hu, X., Fan, K., Zhang, N., Shang, L., Deng, Y., … & Jiang, Z. (2023). Emergence of Corona Is Independent of the Four Whorls of Floral Organs in *Narcissus tazetta*. Plants, 12(7), 1458. 10.3390/plants12071458.

Mazrou, R. M., Hassan, S., Yang, M., & Hassan, F. A. (2022). Melatonin preserves the postharvest quality of cut roses through enhancing the antioxidant system. Plants, 11(20), 2713. 10.3390/plants11202713.

Naing, A. H., Win, N. M., Kyu, S. Y., Kang, I. K., & Kim, C. K. (2022). Current progress in application of 1-methylcyclopropene to improve postharvest quality of cut flowers. Horticultural Plant Journal, 8(6), 676–688. 10.1016/j.hpj.2021.11.014.

Parklak, W., Ounjaijean, S., Kulprachakarn, K., & Boonyapranai, K. (2023). In Vitro α-Amylase and α-Glucosidase Inhibitory Effects, Antioxidant Activities, and Lutein Content of Nine Different Cultivars of Marigold Flowers (*Tagetes* spp.). Molecules, 28(8), 3314. 10.3390/molecules28083314.

Rabiza-Świder, J., Skutnik, E., & Jędrzejuk, A. (2019). The effect of a sugar-containing preservative on senescence-related processes in cut clematis flowers. Notulae Botanicae Horti Agrobotanici Cluj-Napoca, 47(2), 432–440.

Rabiza-Świder, J., Skutnik, E., Jędrzejuk, A., & Łukaszewska, A. (2020a). Postharvest treatments improve quality of cut peony flowers. Agronomy, 10(10), 1583. 10.3390/agronomy10101583.

Rabiza-Świder, J., Skutnik, E., Jędrzejuk, A., & Rochala-Wojciechowska, J. (2020b). Nanosilver and sucrose delay the senescence of cut snapdragon flowers. Postharvest Biology and Technology, 165, 111165. 10.1016/j.postharvbio.2020.111165.

Rabiza-Świder, J., Skutnik, E., Jędrzejuk, A., & Sochacki, D. (2020c). Effect of preservatives on senescence of cut daffodil (*Narcissus* L.) flowers. The Journal of Horticultural Science and Biotechnology, 95(3), 331–340. 10.1080/14620316.2019.1679042.

Skutnik, E., Rabiza-Świder, J., Jędrzejuk, A., & Łukaszewska, A. (2020). The effect of the long-term cold storage and preservatives on senescence of cut herbaceous peony flowers. Agronomy, 10(11), 1631. 10.3390/agronomy10111631.

Spricigo, P. C., Pilon, L., Trento, J. P., de Moura, M. R., Bonfim, K. S., Mitsuyuki, M. C., & Ferreira, M. D. (2021). Nano-chitosan as an antimicrobial agent in preservative solutions for cut flowers. Journal of Chemical Technology & Biotechnology, 96(8), 2168–2175. https://onlinelibrary.wiley.com/doi/10.1002/jctb.6766.

Sun, J., Gu, J., Zeng, J., Han, S., Song, A., Chen, F., … & Chen, S. (2013). Changes in leaf morphology, antioxidant activity and photosynthesis capacity in two different drought-tolerant cultivars of chrysanthemum during and after water stress. Scientia Horticulturae, 161, 249–258. 10.1016/j.scienta.2013.07.015.

Sun, X., Qin, M., Yu, Q., Huang, Z., Xiao, Y., Li, Y., … & Gao, J. (2021). Molecular understanding of postharvest flower opening and senescence. Molecular Horticulture, 1(1), 7. 10.1186/s43897-021-00015-8.

Tarakemeh, A., Azizi, M., Rowshan, V., Salehi, H., Spina, R., Dupire, F., … & Laurain-Mattar, D. (2019). Screening of Amaryllidaceae alkaloids in bulbs and tissue cultures of *Narcissus papyraceus* and four varieties of *N. tazetta*. Journal of Pharmaceutical and Biomedical Analysis, 172, 230–237.

van Doorn, W. G., Sinz, A., & Tomassen, M. M. (2004). Daffodil flowers delay senescence in cut Iris flowers. Phytochemistry, 65(5), 571–577. 10.1016/j.phytochem.2003.12.008.

Verdonk, J. C., van Ieperen, W., Carvalho, D. R., van Geest, G., & Schouten, R. E. (2023). Effect of preharvest conditions on cut-flower quality. Frontiers in Plant Science, 14, 1281456. 10.3389/fpls.2023.1281456.

Vijayakumar, S., Singh, Sh., Pandiyaraj, P., Sujayasree, O. J. (2021). “Post-harvest handling of cut flowers,” in Trends & Prospects in Post-Harvest Management of Horticultural Crops (Today & Tomorrow’s Printers and Publishers, New Delhi - 110 002, India), 419–446.

Waters, M. T., Tiley, A. M., Kramer, E. M., Meerow, A. W., Langdale, J. A., & Scotland, R. W. (2013). The corona of the daffodil *Narcissus bulbocodium* shares stamen-like identity and is distinct from the orthodox floral whorls. The Plant Journal, 74(4), 615–625. doi:10.1111/tpj.12150.

Woltering, E. J., & Paillart, M. J. (2018). Effect of cold storage on stomatal functionality, water relations and flower performance in cut roses. Postharvest Biology and Technology, 136, 66–73. 10.1016/j.postharvbio.2017.10.009.

Zanganeh, R., Jamei, R., & Rahmani, F. (2019). Modulation of growth and oxidative stress by seed priming with salicylic acid in *Zea mays* L. under lead stress. Journal of Plant Interactions, 14(1), 369–375. 10.1080/17429145.2019.1629032.

Zhao, D., Cheng, M., Tang, W., Liu, D., Zhou, S., Meng, J., & Tao, J. (2018). Nano-silver modifies the vase life of cut herbaceous peony (*Paeonia lactiflora* Pall.) flowers. Protoplasma, 255, 1001–1013. 10.1007/s00709-018-1209-1.

Zou, J., Wang, G., Ji, J., Wang, J., Wu, H., Ou, Y., & Li, B. (2017). Transcriptional, physiological and cytological analysis validated the roles of some key genes linked Cd stress in *Salix matsudana* Koidz. Environmental and Experimental Botany, 134, 116–129.

